# Interferon-responsive genes are targeted during the establishment of human cytomegalovirus latency

**DOI:** 10.1101/791673

**Authors:** Elizabeth G. Elder, Benjamin A. Krishna, James Williamson, Eleanor Y. Lim, Emma Poole, George X. Sedikides, Mark Wills, Christine M. O’Connor, Paul J. Lehner, John Sinclair

## Abstract

Human cytomegalovirus (HCMV) latency is an active process which remodels the latently infected cell to optimise latent carriage and reactivation. This is achieved, in part, through the expression of viral genes, including the G-protein coupled receptor US28. Here, we use an unbiased proteomic screen to assess changes in host proteins induced by US28, revealing that interferon-inducible genes are downregulated by US28. We validate that MHC Class II and two PYHIN proteins, MNDA and IFI16, are downregulated during experimental latency in primary human CD14^+^ monocytes. We find that IFI16 is targeted rapidly during the establishment of latency in a US28-dependent manner, but only in undifferentiated myeloid cells, a natural site of latent carriage. Finally, by overexpressing IFI16, we show that IFI16 can activate the viral major immediate early promoter and immediate early gene expression during latency via NF-κB, a function which explains why downregulation of IFI16 during latency is advantageous for the virus.

**Importance:** Human cytomegalovirus (HCMV) is a ubiquitous herpesvirus which infects 50-100% of humans worldwide. HCMV causes a lifelong subclinical infection in immunocompetent individuals, but is a serious cause of mortality and morbidity in the immunocompromised and in neonates. In particular, reactivation of HCMV in the transplant setting is a major cause of transplant failure and related disease. Therefore, a molecular understanding of HCMV latency and reactivation could provide insights into potential ways to target the latent viral reservoir in at-risk patient populations.

## Introduction

Lifelong persistence of Human cytomegalovirus (HCMV) is underpinned by viral latency and reactivation. Following primary infection, the ubiquitous betaherpesvirus HCMV establishes latency in cell types including early myeloid lineage cells ^1–4^. Viral genome is maintained in these cells in the relative absence of immediate early (IE) gene expression or production of infectious virions. Reactivation of HCMV is associated with differentiation of myeloid lineage cells to mature dendritic cells and macrophages; as such, reactivation events are thought to occur sporadically throughout the lifetime of the host ^5–8^. In immunocompetent individuals, both primary infection and reactivation events are well-controlled by a broad and robust immune response ^9^. However, HCMV reactivation is a major cause of morbidity and mortality in immunocompromised patients, including stem cell and organ transplant recipients ^10,11^.

A key hallmark of latency is the relative suppression of IE gene expression ^1,2,12–14^, which is controlled by the major immediate early promoter ^15–17^, and the subsequent lack of infectious virion production. The establishment of latency via MIEP repression in early myeloid lineage cells requires both host and viral factors ^18^. One viral factor that suppresses MIEP activity is the G-protein coupled receptor (GPCR) US28, a virally encoded chemokine receptor homologue, which is expressed *de novo* during latency as well as being delivered to cells with the incoming virion ^19–24^ and this incoming viral US28 is functional ^25^. US28 modulates the signalling pathways of early myeloid cells; it attenuates MAP kinases, NF-κB, and c-fos, whilst activating STAT3 and iNOS ^21,23,25^. All of these contribute to the repression of MIEP activity. This US28-mediated signalling is so critical to latency that US28-deleted viruses, or the loss of G-protein coupling by the US28 mutant R129A, result in lytic infection of undifferentiated myeloid cells ^19,21,23,25^. Furthermore, when examined, these US28-mediated effects on cell signalling did not occur during lytic infection or in permissive cells ^21^, implying that US28 represses the MIEP during latency but does not impair reactivation following cellular differentiation. This is reflective of the cell type-specific nature of US28-mediated signalling ^21,26^.

Since US28 can modulate all these pathways and control the MIEP, we hypothesised that US28 would also cause changes in host protein expression. Here, we perform a proteomic screen comparing host cell protein abundance in myelomonocytic THP-1 cells expressing wild-type (WT) US28 or the US28-R129A signalling mutant. We find that the expression of many host proteins are decreased in the presence of US28-WT compared with the US28-R129A mutant and a large proportion of these proteins are interferon inducible. In particular, the two Pyrin and HIN domain (PYHIN) family proteins, Myeloid Cell Nuclear Differentiation Antigen (MNDA) and Gamma-Interferon-Inducible Protein 16 (IFI16), are downregulated by US28, as well as MHC Class II proteins. IFI16 is associated with the nuclear sensing of herpesvirus DNA ^27–31^ and control of herpesvirus gene expression ^32–39^, and also represses viral transcription during HIV latency ^40^, but the effects of IFI16 on HCMV latent infection is unknown. Downregulation of HLA-DR/MHC Class II is important for the evasion of CD4^+^ T cell responses to latently infected myeloid cells ^41^, while antiviral effects of MNDA have yet to be reported.

We have validated the downregulation of IFI16, MNDA, and HLA-DR in US28-expressing THP-1 cells and during experimental latency in primary CD14^+^ monocytes. We find that HCMV downregulates IFI16 within the first 24 hours of infection of myeloid cells in a US28-dependent manner, but this effect is lost in differentiated dendritic cells. We propose that downregulation of IFI16 is beneficial to the establishment of latency because overexpression of IFI16 drives MIEP activity and IE gene expression via NF-κB. By targeting the downregulation of IFI16, US28 actively promotes the establishment of latency in early myeloid lineage cells.

## Materials and Methods

### Cells

All cells were maintained at 37 °C in a 5% CO_2_ atmosphere. THP-1 cells (ECACC 88081201) were cultured in RPMI-1640 media (Sigma) supplemented with 10% heat-inactivated fetal bovine serum (FBS; PAN Biotech), 100 U/mL penicillin and 100 µg/mL streptomycin (Sigma), and 0.05 mM 2-mercaptoethanol (Gibco). Kasumi-3 cells (ATCC® CRL-2725) were cultured in RPMI-1640 media (Sigma) supplemented with 20% heat-inactivated fetal bovine serum (FBS; PAN Biotech), 100 U/mL penicillin and 100 µg/mL streptomycin (Sigma). During infections, THP-1 and Kasumi-3 cells were cultured in a low-serum (1%) version of this media for a minimum of 24 hours prior to inoculation, and maintained in this low-serum media throughout the infection. MIEP-eGFP reporter THP-1 cells ^42^ were a gift from M Van Loock, Johnson & Johnson. RPE-1 cells (ATCC® CRL-4000™) and Human foreskin fibroblasts (Hff1; ATCC® SCRC-1041™) were maintained in DMEM (Sigma) supplemented with 10% heat-inactivated FBS and 100 U/mL penicillin and 100 µg/mL streptomycin. 293T cells (ECACC 12022001) were maintained in DMEM (Sigma) supplemented with 10% heat-inactivated FBS but without penicillin or streptomycin. Phorbol 12-myristate 13-acetate (PMA, Sigma) was used to induce myeloid cell differentiation at a concentration of 20 ng/mL.

Primary CD14^+^ monocytes were isolated from apheresis cones (NHS Blood & Transplant Service) or from peripheral blood taken from healthy volunteers as previously described ^43^. Briefly, CD14^+^ monocytes were isolated from total PBMC by magnetic-activated cell sorting (MACS) using CD14^+^ microbeads (Miltenyi Biotech). The monocytes were plated on tissue culture dishes (Corning) or slides (Ibidi), or kept in suspension in X-Vivo 15 media (Lonza) supplemented with 2 mM L-glutamine. Mature dendritic cells were generated by treating CD14^+^ monocytes with granulocyte-macrophage colony-stimulating factor (GM-CSF, Miltenyi, 1000U/mL) and interleukin-4 (IL-4, Miltenyi, 1000U/mL) for 5 days before addition of lipopolysaccharide (LPS, Invivogen, 50 ng/mL) for 2 further days.

Primary human CD34^+^ hematopoietic progenitor cells, isolated from adult bone marrow, were purchased from STEMCELL Technologies and cultured in X-Vivo 15 media (Lonza).

### Generation of lentiviruses and retroviruses

The lentiviral vectors encoding US28 from the VHL/E strain of HCMV have been described previously ^21^; US28 is expressed in these vectors via the SFFV promoter. The lentiviral vectors pHRSIN UbEm and pHRsin SV40blast were a kind gift from D. van den Boomen, University of Cambridge, and were based upon a previously published lentiviral system^44,45^. Briefly, expression of the gene of interest is also driven by the Spleen Focus-Forming Virus (SFFV) promoter, and the selectable markers Emerald and blasticidin resistance are driven by the Ubiquitin promoter (UbEm) and the SV40 promoter (SV40blast), respectively. The sequence encoding US28 from the VHL/E strain of HCMV was cloned into pHRSIN UbEm using the EcoR1 and Spe1 sites using the following primers: US28 FW 5’ GCACGAATTCCATATGACGCCGACGACGAC AND RV 5’ CTGCACTAGTTTACGGTATAATTTGTGAGAC. The sequence encoding IFI16 was cloned into pHRsin SV40blast using the BamHI and NotI sites using the following primers: IFI16 FW 5’ GATTGCGGCCGCATGGGAAAAAAATACAAGAACATTGTTC and RV 5’ GATCGGATCCTTAGAAGAAAAAGTCTGGTGAAGTTTC.

The sequence encoding US28-3XFLAG was cloned from TB40/E*mCherry*-US28-3XFLAG into the retroviral plasmid pBABE eGFP (a gift from Debu Chakravarti (Addgene plasmid #36999)) as described previously^25^. The Q5 site directed mutagenesis kit (New England Biotech) was used to generate the US28-R129A mutant of this vector, which was verified by Sanger Sequencing. Expression of US28 in these vectors is driven by the long terminal repeat and partial gag.

Generation of VSV-G pseudotyped lentiviral particles was conducted generally in line with the Broad Institute Protocols. Five hundred thousand 293T cells were transfected with 1250 ng of lentiviral expression vector, 625 ng of lentiviral packaging vector psPAX and 625 ng of envelope vector pMD.2G (both gifts from S. Karniely, Weizmann Institute, Israel) using transfection reagent FuGene6 (Promega) according to the manufacturer’s instructions. For generation of VSV-G pseudotyped retrovirus particles, 1250 ng of the murine leukemia virus retroviral packaging vector KB4 ^46^ (a gift from H. Groom, University of Cambridge) was transfected along with 625 ng pMD.2G and 1250 ng retroviral expression vector.

### Lentiviral and retroviral transduction

Supernatants from transfected 293T cells were harvested at 36 and 60 hours post transfection, filtered through a 0.45 µm syringe filter, and used to transduce THP-1 cells in the presence of 2 μg/mL polybrene. When necessary, lentiviral titres were determined by in-house p24 enzyme-linked immunosorbent assay (ELISA). For transduction with puromycin-resistance vectors, puromycin (2 μg/mL, Sigma) was added to media and refreshed every 2-5 days until all control non-transduced THP-1 cells were dead. Similarly, where blasticidin-resistance vectors were used, blasticidin (1 μg/mL, Invivogen) was added to media. Emerald positive cells were sorted using a BD FACSAriaIII instrument.

### Human cytomegaloviruses

Infection of monocytes and THP-1 cells were carried out at a multiplicity of infection (MOI) of 3 as determined by titration on RPE-1 cells. TB40/E BAC4 strains were propagated in RPE-1 cells by seeding 50% confluent T175 flasks with virus at an MOI of 0.1. Spread of virus was monitored for 2-6 weeks following inoculation by fluorescence microscopy, and infected monolayers were subcultured twice during this period. When cells were 95-100% infected, supernatants were harvested on three occasions spaced over 7 days and stored at −80°C. In the final harvest, the monolayer was scraped and also stored at −80°C. After thawing the virus-containing media, cell debris was pelleted by centrifugation at 1500 x *g* for 20 minutes at RT. Then, the clarified supernatant was concentrated by high speed centrifugation at 14500 x *g* for 2 hours at 18°C. Virus-containing pellets were then resuspended in X-vivo-15 media in aliquots at −80°C.

TB40/E*mCherry*-US28-3XFLAG and TB40/E*mCherry*-US28Δ have been described previously ^19^. TB40/E*gfp* ^47^ and TB40/E BAC4 SV40 mCherry IE2-2A-GFP^48^ were kind gifts from E.A. Murphy, SUNY Upstate Medical University. TB40/E BAC4 IE2-eYFP has been described previously ^49,50^. TB40/E BAC4 GATA2mCherry has been described previously^51^. TB40/E with deleted NF-κB sites in the MIEP at positions −94, −157, −262 and −413, and the revertant virus, were a kind gift from Jeffery Meier and Ming Li (University of Iowa, United States), and have been described previously^52^.

UV-inactivation of virus was performed by placing a 100 μL aliquot of virus in one well of a 24-well plate and placing this within 10cm of a UV germicidal (254 nm) lamp for 15 minutes, which routinely results in no detectable IE gene expression upon infection of Hff1 cells.

### Immunofluorescence staining and image analysis

Cells were fixed with 2% paraformaldehyde for 15 minutes and permeablised with 0.1% Triton-X100 for 10 minutes at RT. Blocking and antibody incubations were performed in phosphate buffered saline (PBS) with 1% bovine serum albumin and 5% normal goat serum. Antibodies used: anti-IFI16 (Santa Cruz sc-8023, 1:100), anti-FLAG (Sigma F1804, 1:1000), anti-MNDA (Cell Signaling Technology 3329, 1:100), anti-IE (Argene 11-003, 1:1000 or directly conjugated to FITC, 1:100), anti-GFP (directly conjugated to FITC, Abcam ab6662, 1:200), anti-mCherry (Abcam ab167453, 1:500), anti-HLA DR (conjugated to Brilliant Violet 421, Biolegend Clone L423 or Abcam ab92511 1:100). Cells were imaged with a widefield Nikon TE200 microscope and images were processed using ImageJ. For contingency analyses of IFI16 expression during experimental latency, cells were assigned as ‘IFI16 positive/negative’, and ‘infected/uninfected’ and then counted. These results were then analysed using Fischer’s Exact statistical test for significance. For analysis of signal intensity, nuclear stained images were used to create a mask from which intensity values of the corresponding IFI16/MNDA stained image were derived using the Analyze Particles feature of ImageJ. Cells were assigned as infected or uninfected based on signal from the GFP/mCherry stain. The average signal intensity of uninfected cells was used to normalise the signal intensity in order to compare different fields of view.

### Inhibitors

The c-fos inhibitor T5524 was purchased from Cayman Chemical, solubilised in DMSO and used at 10 μM. The Janus kinase inhibitor Ruxolitinib was purchased from Cell Guidance Systems, solubilised in DMSO and used at 5 μM. The IKKα inhibitor/NF-κB pathway inhibitor BAY11-7082 was purchased from Santa Cruz, solubilised in DMSO, and used at a concentration of 5 μM.

### Western blotting

Except for US28 blots, cells were lysed directly in Laemmli Buffer and separated by SDS-PAGE. Following transfer to nitrocellulose, the membrane was blocked in 5% milk in tris buffered saline (TBS) with 0.1% Tween-20. Antibodies used: anti-IFI16 (Santa Cruz sc-8023, 1:500), anti-MNDA (Cell Signaling Technology 3329, 1:250), anti-STAT1 (Cell Signaling Technology 14994, 1:1000), anti-phosphoSTAT1 Tyr701 (Cell Signaling Technology 9167, 1:1000, anti-beta actin (Abcam ab6276, 1:5000). For US28 blots, cells were pelleted, washed once in ice cold PBS, then lysed in native lysis buffer (25 mM Tris HCl pH 7.4, 150 mM NaCl, 1 mM EDTA, 1% NP40, 5% glycerol, plus protease inhibitors) for 30 minutes, vortexing every 10 minutes. After the addition of non-reducing Laemmli buffer, samples were heated at 42°C for 10 minutes and then separated by SDS-PAGE. Polyvinylidene difluoride membranes were used for transfer, and blocked membranes were incubated with the rabbit anti-US28 antibody^53^ (a gift from M. Smit, Vrije University) at 1:1000 dilution. To quantify western blots, the Analyze Gels feature of Image J was used to plot the band intensities. Signal for genes of interest were then normalised to US28-R129A cells and then normalised to either the relative amounts of actin or STAT1, as described in the figure legends.

### Flow cytometry

Transduced THP-1 cells and MIEP-reporter THP-1 cells were analysed on a BD Accuri Instrument. Live cells were gated using forward and side scatter. Paraformaldehyde-fixed cells were stained using anti-HLA-DR APC conjugate (Biolegend, Clone L243, 1:50). Latently infected CD14^+^ monocytes were fixed with 1% paraformaldehyde and stained using anti-HLA-DR Pacific blue conjugate (Biolegend, Clone L243, 1:50) and anti-HLA-A,B,C, PE-Cy7 conjugate (Biolegend, Clone W6/32, 1:50), before analysis on the Nxt Attune Instrument (Thermo Fisher).

### RNA extraction, reverse transcription and quantitative PCR

RNA was extracted and purified using Direct-Zol RNA MiniPrep kit (Zymo Research) according to the manufacturer’s instructions. A total of 5 ng of purified RNA was used in RT-qPCR reactions, performed using QuantiTect SYBR® Green RT-PCR Kit reagents (Qiagen) on a StepOne Real-Time PCR instrument (Applied Biosystems). TATA-box binding protein (TBP) was used as a reference gene and fold changes were analysed by the 2-ΔΔCt method. Reverse transcription was performed using the Qiagen QuantiTect Reverse Transcription kit, and then cDNA was used in qPCR analysis using New England Biotech LUNA SYBR Green qPCR reagents using TBP or GAPDH as a reference gene. Primers used: US28 FW: AATCGTTGCGGTGTCTCAGT; US28 RV: TGGTACGGCAGCCAAAAGAT; MNDA FW: GGAAGAAGCATCCATTAAGG; MNDA RV: GTTTGTCTAGACAGGCAAC; IFI16 FW: CTGCACCCTCCACAAG; IFI16 RV: CCATGGCTGTGGACATG; TBP FW: CGGCTGTTTAACTTCGCTTC; TBP RV: CACACGCCAAGAAACAGTGA; HLA-DRA FW: TGTAAGGCACATGGAGGTGA; HLA-DRA RV: ATAGGGCTGGAAAATGCTGA; IE72 FW: GTCCTGACAGAACTCGTCAAA; IE72 RV: TAAAGGCGCCAGTGAATTTTTCTTC; GAPDH FW TGCACCACCAACTGCTTAGC, GAPDH RV: GGCATGGACTGTGGTCATGAG.

### Sequence alignment

US28 sequences were exported from GenBank (TB40/E BAC4 EF999921.1, AD169 FJ527563.1, Merlin NC_006273.2, VHL/E MK425187.1) or from sequenced plasmids, and aligned using Clustal Omega^54^ https://www.ebi.ac.uk/Tools/msa/clustalo/ and the output format MView^55^.

### Cell lysis, digestion and cleanup for proteomic analysis

Cells were harvested by centrifugation and washed 3x in cold phosphate-buffered saline (PBS) before finally pelleting into a low adhesion microfuge tube. Cell pellets were lysed in 2%/50 mM Triethylamminium bicarbonate (TEAB) pH 8.5. Samples were quantified by BCA assay and 50μg of each sample was taken and adjusted to the same volume with lysis buffer. Reduction and alkylation was achieved by addition of 10 mM Tris (2-carboxyethyl)phosphine(TCEP) and 20 mM iodoacetamide for 20mins at room temperature in the dark followed by quenching with 10mM DTT. Samples were further purified and digested using a modified Filtered Aided Sample Prep (FASP) protocol. Briefly, samples were brought to 500 μL volume with 8 M urea/TEAB and loaded onto a 30kDa molecular weight cut-off (MWCO) ultrafiltration device. Samples were then washed 3 times with 8 M urea/TEAB followed by 3 times with 0.5% deoxycholate (SDC) /50mM TEAB. Samples were finally resuspended in ~50 μL of SDC /TEAB containing 1 μg trypsin and incubated overnight at 37°C. After digestion samples were recovered from the filter device by centrifugation, acidified to precipitate SDC and cleaned up by two-phase partitioning into 2x volumes of ethyl acetate (repeated twice) before drying in a vacuum centrifuge.

### TMT Labelling

Samples were resuspended in 20 μL 100 mM TEAB and to each tube 0.2 μg of a unique tandem mass tag (TMT) label for each sample was added in 8.5 µL acetonitrile (ACN) and incubated for 1 h at room temperature. TMT reactions were quenched by addition of 3 µL of 200 mM ammonium formate, pooled and dried in a vacuum centrifuge. The sample was then resuspended in 800µL 0.1% trifluoroacetic acid (TFA) and acidified to ~pH 2 with formic acid (FA) before performing a C18-solid phase extraction (C18-SPE) using a Sep-Pak cartridge (Waters) attached to a vacuum manifold. C18 eluate was dried in a vacuum centrifuge and resuspended in 40 µL 200 mM ammonium formate, pH 10.

### High pH Revered Phase Fractionation

Sample was injected onto an Ultimate 3000 RSLC UHPLC system (Thermo Fisher Scientific) equipped with a 2.1 i.d. x25cm, 1.7µm particle Kinetix Evo C18 column (Phenomenex). Mobile phase consisted of A: 3% ACN, B: ACN and C: 200mM ammonium formate pH 10. Isocratic conditions were 90% A/10%C and C was maintained at 10% throughout the gradient elution. Separations were carried out at 45°C. After loading at 200 µL/min for 5 min and ramping the flow rate to 400 µL/min over 5 min, the gradient elution proceeded as follows: 0-19% B over 10 minutes (curve 3), 19-34% B over 14.25mins (curve 5), 34-50% B over 8.75mins (curve 5), followed by a 10 min wash at 90% B. UV absorbance was monitored at 280 nm and 15 s fractions were collected into 96 well microplates using the integrated fraction collector. Peptide containing fractions were then orthogonally recombined into 24 fractions and dried in a vacuum centrifuge and resuspended in 10 µL 5% DMSO 0.5% TFA for analysis.

### LC-MS analysis

All samples were injected onto an Ultimate 3000 RSLC nano UHPLC equipped with a 300 µm i.d. x 5 mm Acclaim PepMap µ-Precolumn (Thermo Fisher Scientific) and a 75 µm i.d. x50c m 2.1 µm particle Acclaim PepMap RSLC analytical column. Loading solvent was 0.1% TFA, analytical solvent A: 0.1% FA and B: ACN+0.1% FA. All separations are carried out at 55°C. Samples were loaded at 10 µL/min for 5 min in loading solvent before beginning the analytical gradient. For high pH reverse phase (RP) fractions, a gradient of 3-5.6% B over 4 min, 5.6 – 32% B over 162 min, followed by a 5 min wash at 80% B and a 5 min wash at 90% B and equilibration at 3% B for 5 min. During the gradient the Orbitrap Fusion mass spectrometer (Thermo Fisher Scientific) was set to acquire spectra according to the settings given in supplementary file S6 “MS Settings”.

### Data Processing

All raw files were searched by Mascot within Proteome Discoverer 2.1 (Thermo Fisher Scientific) against the Swissprot human database and a database of common contaminants.

The search parameters were as follows. Enzyme: Trypsin. MS1 tol: 10 ppm. MS2 tol: 0.6 Da. Fixed modifications: Carbamidomethyl cysteine, TMT peptide N-termini and lysine. Variable modification oxidised methionine. MS3 reporter ion tol: 20 ppm, most confident centroid. Mascot Percolator was used to calculate peptide-spectrum match false discovery rate (PSM FDR).

Search results were further processed and filtered as follows: Peptides below a percolator FDR of 0.01% and proteins below the 0.01% protein FDR (calculated from a built-in decoy database search) were rejected. Protein groups were then generated using the strict parsimony principle. Peptides both unique and razor with a co-isolation threshold of 50 and an average signal-to-noise (s/n) threshold of 10 were used for quantification and a normalisation of these values to the total peptide amount in each channel was applied. Instances where a protein was identified but not quantified in all channels were rejected from further analysis. “Scaled” abundances of proteins provided by Proteome Discoverer were used to derive ratios of abundance. Q values of significance between groups were calculated by Benjamini-Hochberg correction of p values generated using the moderated T-test LIMMA within the R environment.

## Results

### Proteomic analysis reveals US28-induced changes in host proteins in myeloid cells

US28-mediated signalling is critical to latency: US28-deleted viruses, or the loss of G-protein coupling by the US28 mutant R129A, result in lytic infection of undifferentiated myeloid cells ^19,21,23,25^. Similarly, infection of US28-WT-expressing THP-1 cells with US28-deletion viruses leads to complementation and the establishment of latency; both the US28-R129A and empty vector cell lines fail to establish latent infection under these conditions ^21^. Therefore, to understand how US28-WT alters the host cell environment to support latency, we analysed the total proteomes of myelomonocytic THP-1 cell lines which express either US28-WT (sequence derived from the VHL/E strain of HCMV), signalling mutant US28-R129A, or the empty vector (EV) control (Figures 1A, 1B, 1C, File S5).

**Figure 1.**
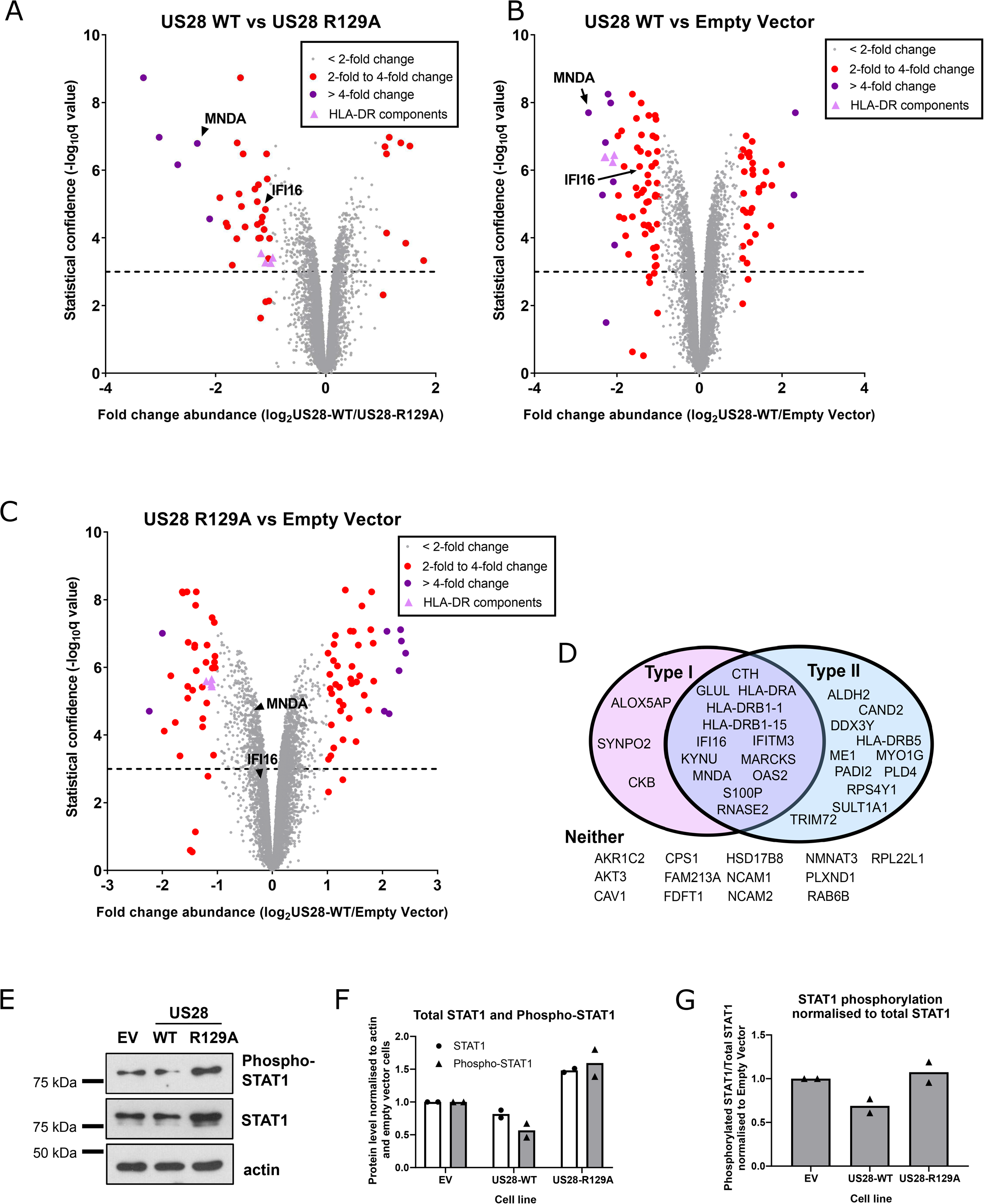
US28 induces changes in the host proteome of THP-1 cells. A, B, C) THP-1 cells expressing empty vector, US28-WT, and US28-R129A were subject to total cell proteomic analysis using a TMT labelling approach as described in Materials and Methods. Each dot represents one human protein and is shown in grey if its abundance changes by a factor of less than 2, in red if between 2- and 4-fold, and in purple if greater than 4-fold. The exception is components of the HLA-DR complex which are represented by pink triangles. The dotted line represents a significance threshold of q = 0.001; q < 0.001 is considered significant. Q values of significance between groups were calculated by Benjamini-Hochberg correction of p values generated using the moderated T-test LIMMA within the R environment. Comparison of A) US28-WT and US28-R129A, B) US28-WT and empty vector, and C) US28-R129A and empty vector. In each case, the relative abundance of human proteins MNDA and IFI16 is marked with an arrow. D) Analysis of the top 40-downregulated proteins identified in (A). After filtering for changes with a q <0.001, the gene names were entered in the Interferome database. Proteins that are induced by Type I and/or Type II interferon are depicted in the Venn diagram. Proteins we identified that are not interferon-inducible are listed below (‘Neither’). E) Lysates from THP-1 cells expressing Empty Vector (EV), US28-WT and US28-R129A were assessed by western blot for phospho-STAT1 (Tyr701), total STAT1, and beta-actin (loading control). F and G) Quantification of STAT1 and phospho-STAT1 band intensity from two western blots from two independent samples of transduced THP-1 cells. F) Quantification of the indicated protein levels normalised to actin. G) Quantification of phospho-STAT1 levels relative to total STAT1.

By using these three cell lines, we were able to identify host cell proteins modulated by US28 expression in a G-protein signalling dependent and independent manner. Changes in host protein abundance common between US28-WT and US28-R129A when compared to empty vector (Figure 1B and 1C) represent signalling independent changes, and these include CD44 and CD82 proteins, which are each downregulated by both sets of US28-expressing cells.

While we do not rule out that signalling independent changes in myeloid cells driven by US28 may be important for HCMV latency, G-protein dependent signalling is absolutely required for latency, and therefore we were particularly interested in the direct comparison of host protein abundances in THP-1 cells expressing US28-WT and US28-R129A (Figure 1A). This comparison reveals 42 host proteins whose expression is two-fold or more increased or decreased by US28-WT, and our analyses focussed on these signalling-dependent changes.

One remarkable feature of many of the most downregulated proteins in Figure 1A is that they are interferon-inducible (Figure 1D, Figure S1A). According to the Interferome database (v2.01, www.interferome.org) ^56^, two-thirds (27/40) of the most downregulated proteins (fold change 1.86-fold or higher) we identified are Type I or Type II interferon-inducible (Figure 1D). In contrast, of the 40 proteins which showed no changes (fold change = 0) in abundance between US28-WT and US28-R129A, 12/40 (30%) were included in the Interferome database (File S5). Importantly, overexpression of the multipass membrane US28 proteins did not lead to induction of the unfolded protein response or other endoplasmic stress related genes (Figure S1B), suggesting that the changes identified in the screen are not general effects of protein overexpression.

Since STAT1 phosphorylation is common to both the Type I and Type II interferon signalling pathways ^57^, we examined total STAT1 and phosphorylated STAT1 in US28-WT cells. We found that US28-WT cells had lower overall levels of STAT1 and phosphorylated STAT1 compared to cells transduced with US28-R129A or empty vector control (Figure 1E and 1F). Furthermore, when correcting for total levels of STAT1, we found that US28-WT cells had lower relative levels of phosphorylated STAT1 compared with US28-R129A (Figure 1E and 1G). This could help explain why US28-WT downregulated many interferon-inducible genes in our proteomic screen.

### IFI16, MNDA, and HLA-DR are all downregulated by US28

Several interferon-inducible proteins showing decreased expression were of interest as potentially important targets for US28 during HCMV latency. These included MNDA (9 unique peptides; 5.0-fold downregulated compared to US28-R129A), IFI16 (4 unique peptides; 2.4-fold downregulated compared to US28-R129A), and components of the MHC Class II HLA-DR complex (3-6 unique peptides; between 1.9 and 2.2-fold downregulated compared to US28-R129A). We began by confirming US28-WT-mediated downregulation of these proteins in independently-transduced US28-expressing THP-1 cells. After generating these fresh US28-expressing cell lines, we checked expression levels of US28-WT or US28-R129A by RT-qPCR (Figure S2A) and Western blot (Figure S2B and S2C). RT-qPCR confirmed that IFI16, MNDA, and HLA-DRA transcripts are all downregulated in US28-WT-expressing cells compared to those expressing the signalling mutant R129A (Figure 2A). Subsequently, we confirmed this US28-WT mediated downregulation of IFI16 and MNDA at the protein level by western blot (Figure 2B, 2C, S2D, S2E, S2F) or by flow cytometry for cell surface HLA-DR (Figure 2F and 2G), although both US28-WT and US28-R129A cells responded similarly to IFN-γ stimulation, by upregulating HLA-DR similar levels (Figure 2F). This suggests that, whilst US28 is able to downregulate constitutive HLA-DR expression, it is unable to prevent its induction by IFN-γ. Furthermore, this downregulation was not due to strain-specific effects of US28 as US28-WT also downregulated HLA-DR when the US28 sequence from the TB40/E strain of HCMV was used to transduce THP-1 cells (Figure S3A-D).

**Figure 2.**
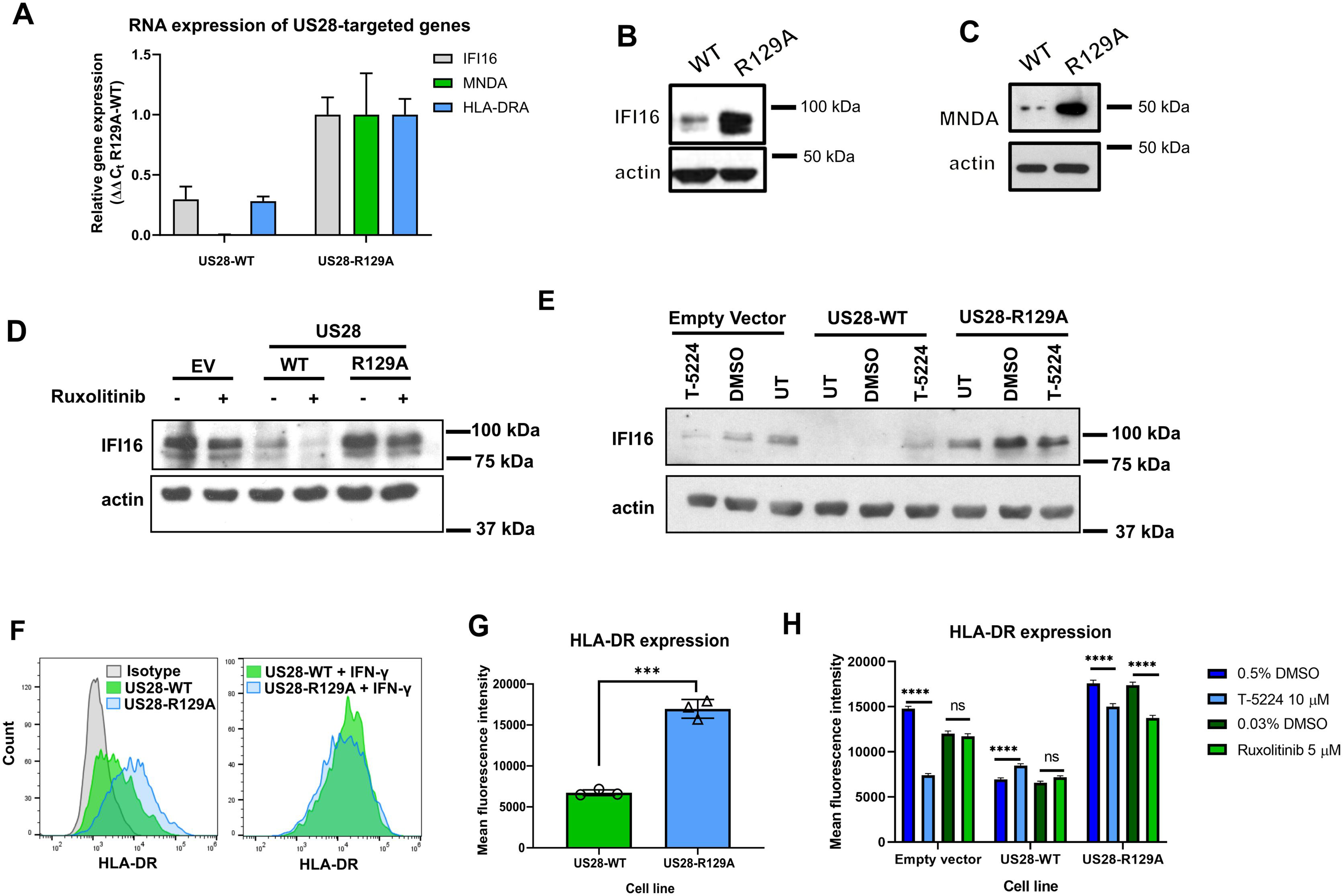
US28-expressing cell lines downregulate IFI16, MNDA, and HLA-DR. A) Relative RNA expression of IFI16, MNDA, and HLA-DR in US28-expressing THP-1 cells. Levels of IFI16, MNDA, HLA-DRA were normalised to TBP and then to US28-R129A using the ΔΔC_t_ method. B and C) Lysates from US28-WT and US28-R129A expressing cells were analysed by western blot for IFI16 (B) and MNDA (C) expression; actin is shown as a loading control. D) Empty vector (EV), US28-WT, and US28-R129A expressing cells were treated with 5 μM ruxolitinib (+), or an equivalent concentration of DMSO (-), for 48 hours, before analysis for IFI16 expression by western blot, using actin as a loading control. E) As D), except cells were treated with T-5224 at 10 μM, DMSO, or left untreated. F) US28-expressing cells were maintained in culture media only (left panel) or treated with 1 ng/mL of IFN-γ (right panel) for 24 hours before staining for cell-surface HLA-DR by flow-cytometry. Staining was performed in triplicate for untreated cells and the mean of these is experiments is presented in G) as mean fluorescence intensity with standard deviation. Statistical analysis was performed by one-way ANOVA; *** P<0.001. H) As D and E), except cells were analysed for cell-surface HLA-DR by flow-cytometry and results are presented as mean fluorescence intensity with 95% confidence intervals. Statistical analysis was performed by two-way ANOVA using Boniferri’s multiple comparison test; ns P>0.01, **** P<0.0001.

Since US28 attenuates c-fos signalling^25^ and STAT1 phosphorylation (Figure 1E, 1F, 1G), and both HLA-DR and IFI16 are fos and STAT1-responsive genes^58–62^, we hypothesised that one or both of these mechanisms is responsible for US28-mediated downregulation of IFI16 and HLA-DR. To test this, we treated empty vector, US28-WT, and US28-R129A expressing THP-1 cells with the Janus Kinase inhibitor Ruxolitinib, or the c-fos inhibitor T-5224. Ruxolitinib partially downregulated IFI16 expression in all three cell types (Figure 2D), but downregulated HLA-DR only in the US28-R129A cell line (Figure 2H), despite a complete block in STAT1 phosphorylation (Figure S2G). The c-fos inhibitor reduced IFI16 and HLA-DR expression in comparison with DMSO controls in empty vector and R129A cell lines, and in the case of empty vector, this drop in expression was almost down to the level in untreated US28-WT cells (Figure 2E and 2H). Taken together, we think it likely that both c-fos and STAT1 attenuation are important for US28-mediated downregulation of IFI16 and HLA-DR. Interestingly, the c-fos inhibitor actually increased IFI16 expression in US28-WT expressing cells. We think this could be due to a basal level of c-fos being required for the expression of a host gene, as yet unidentified, that is need by US28-WT to attenuate multiple signalling pathways; one candidate gene for this is the AP-1 inducible phosphatase DUSP1^63^ which will require further investigation.

### Latent infection of monocytes is associated with the downregulation of IFI16, MNDA, and HLA-DR

Having confirmed key observations from our proteomic data in transduced THP-1 cells overexpressing US28 in isolation, we then sought to determine whether IFI16, MNDA, and HLA-DR are also downregulated in an experimental model of latency in *ex vivo* primary CD14^+^ monocytes where latency-associated expression of US28 is well established. To do this, we infected CD14^+^ monocytes with TB40/E-BAC4 strains of HCMV engineered to express either GFP or mCherry as markers for latent or lytic infection. Firstly, we analysed CD14^+^ monocytes infected with TB40/E SV40 mCherry/IE2-2A-GFP. This virus drives constitutive mCherry expression in all infected cells via the SV40 promoter, but GFP expression is restricted to lytically infected cells as a result of IE2 expression, which is linked to GFP by the self-cleaving peptide 2A. Therefore, we were able to distinguish IE2-positive (lytic) from IE2-negative cells (one hallmark of latency) amongst infected, mCherry positive cells. At four days post infection (d.p.i.), we fixed and immunostained the monocytes for our cellular proteins of interest in mCherry positive, IE2-2A-GFP negative cells (Figure 3A). As a control, we also differentiated monocytes with phorbol 12-myristate 12-acetate (PMA), which drives IE2-2A-GFP expression through differentiation-dependent reactivation. We found that IFI16, MNDA, and HLA-DR were all downregulated in latently infected, mCherry positive but IE2-negative, CD14^+^ monocytes. Importantly, this comparison held true when comparing infected monocytes with uninfected monocytes that had not had contact with viral particles (Figure 3A, S4A).

**Figure 3.**
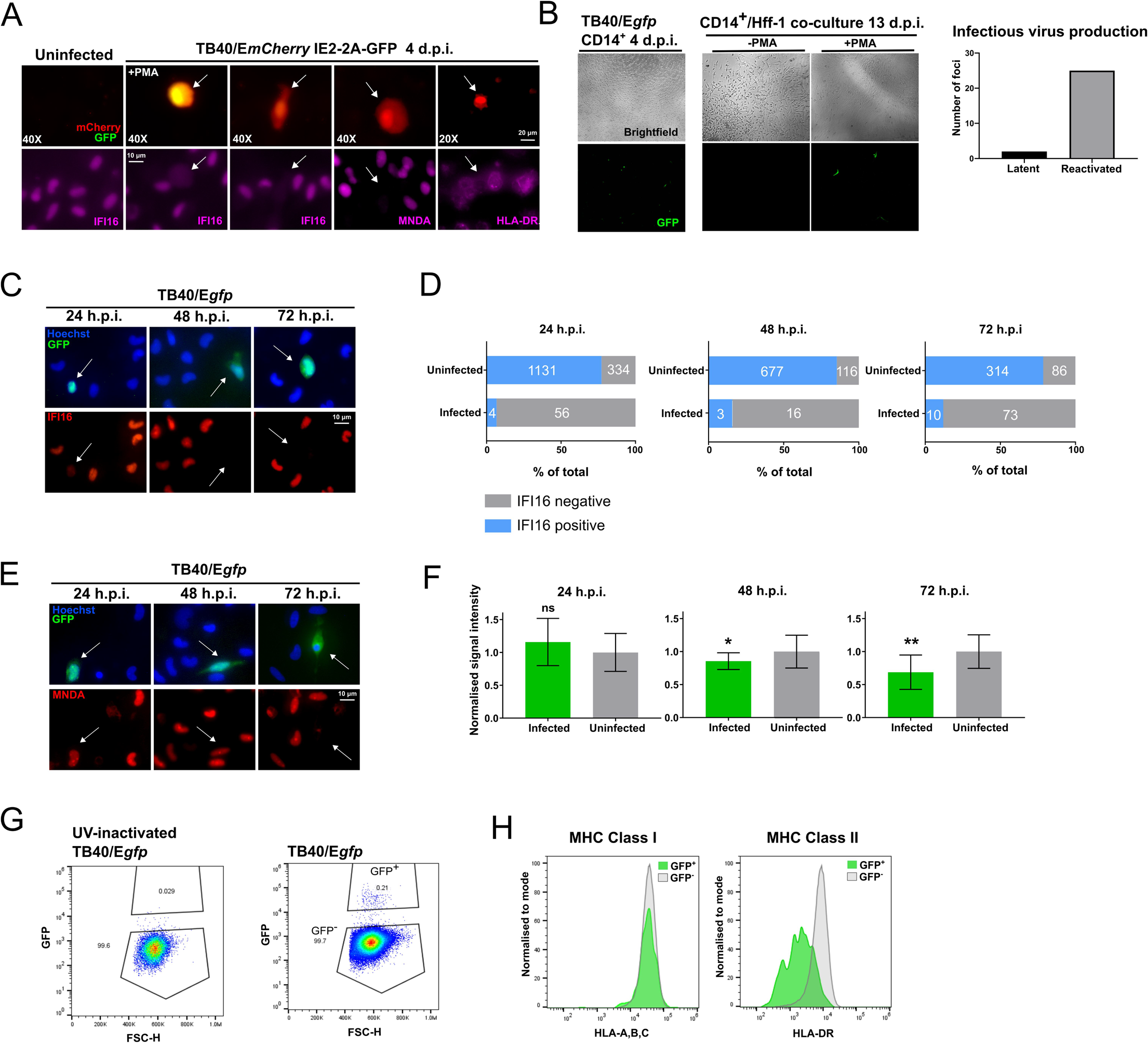
IFI16, MNDA, and HLA-DR are downregulated in latently infected CD14^+^ monocytes. Primary CD14^+^ monocytes were isolated from peripheral blood or apheresis cones as described in Materials and Methods. These cells were then infected using BAC-derived strains of TB40/E. A) CD14^+^ monocytes latently infected with TB40/E SV40-mCherry IE2-2A-GFP stained by immunofluorescence for IFI16, MNDA, or HLA-DR as indicated at four d.p.i and imaged by widefield fluorescence microscopy. Top left image: Uninfected monocytes. Second from the left: Monocytes were treated +PMA (to permit lytic infection). mCherry (red) serves as a marker for infection and GFP (green) denotes expression from the IE2-2A-GFP cassette. Remaining panels: Monocytes were cultured in the absence of PMA. The absence of green fluorescence results from suppressed expression of the IE2-2A-GFP cassette and scored as IE negative. The magnification is indicated (40X or 20X). White arrows indicate corresponding cells in the upper and lower panels. B) Validation of experimental latency using TB40/E*gfp* virus. CD14^+^ Monocytes were infected and allowed to establish latency for 4 days (left panel, 10X magnification). Citrate wash buffer was used to remove externally bound virions. These latently infected cells were cultured -/+PMA for 3 days, and at 7 d.p.i, Hff-1 cells were added to the culture to demonstrate production of infectious virions. Transfer of virus to Hff-1 was monitored by fluorescence microscopy up to 13 d.p.i., and infected Hff-1 foci were counted and summed across the experiment (three wells of CD14^+^ monocytes per condition, graphed). C) CD14^+^ monocytes infected with TB40/E*gfp* stained by immunofluorescence for IFI16 at 24, 48, and 72 h.p.i. and imaged as before using 60X magnification. D) Quantification of IFI16 positive and negative monocytes in the uninfected and infected populations from two donors per time point. Raw numbers of cells are indicated in white text. Fisher’s exact test indicates a statistically significant difference between uninfected and infected populations for each time point (P<0.0001). E) CD14^+^ monocytes infected with TB40/E*gfp* were stained by immunofluorescence for MNDA at the indicated times and imaged as before using 60X magnification. F) Quantification of the signal intensity from infected monocytes at the indicated time points (n=9,7,10, respectively). MNDA signal intensity in each nucleus was normalised to the average of uninfected monocytes from each field of view. A *t*-test with Welch’s correction was used to determine statistical significance. ns, not significant, *P<0.05, **P<0.01 G) CD14^+^ monocytes infected with TB40/E*gfp* (+/-UV inactivation) were analysed for HLA-ABC and HLA-DR expression at three d.p.i. by flow cytometry. The gating strategy for identifying infected cells (GFP^+^) is shown. H) Histogram showing HLA-ABC and HLA-DR staining in HCMV-uninfected GFP-negative (grey) monocytes, and latently infected GFP positive (green) monocytes.

We then sought to look at expression of these proteins at earlier time points. US28 is a virion-associated protein ^19^, and incoming US28 is reported to have rapid effects on host cells ^25^. We speculated that the downregulation of IFI16, MNDA, and HLA-DR might occur early during the establishment of latency. For these experiments, we used TB40/E*gfp* which marks infected cells with GFP expression via the SV40 promoter and confirmed the establishment of latency in this system by coculture of monocytes with fibroblasts either with or without PMA-induced reactivation (Figure 3B). In our latency system, we found a stark and specific loss of IFI16 in infected monocytes from 24 hours post infection (h.p.i.) (Figure 3C), a phenotype maintained at 48 and 72 h.p.i (Figure 3C) as measured by immunofluorescence. We quantified these observations in several fields of view for each of these three time points, and performed contingency analyses (Fisher’s Exact), which confirmed specific loss of IFI16 in latently infected cells (Figure 3D). Loss of IFI16 was observed in *ex vivo* infected monocytes at these time points in a total of four separate donors with TB40/E*gfp* virus.

We found a partial downregulation of MNDA by 72 h.p.i (Figure 3E and 3F), with a very small downregulation at 48 h.p.i and no effect at 24 h.p.i, suggesting that modulation of MNDA is delayed compared with fellow PYHIN family member, IFI16. We also observed that HLA-DR, but not corresponding MHC Class I HLA-A,B,C, were downregulated at 72 h.p.i specifically in GFP positive, latently infected monocytes (Figure 3G and 3H).Therefore, IFI16, MNDA, and HLA-DR are indeed downregulated at early times during the establishment of latency, with IFI16 showing downregulation within 24 hours of infection.

### The downregulation of IFI16 is dependent on viral US28

Having confirmed that IFI16 is downregulated very early during latent infection of monocytes, we then sought to establish whether this effect is dependent on US28. We predicted this would be the case because of the results of our US28 proteomic screen and the established functionality of incoming virion-associated US28 ^25^. We infected monocytes with either the US28 WTTB40/E*mCherry*-US28-3XFLAG HCMV (US38-3XF), or the corresponding US28 deletion virus TB40/E*mCherry*-US28Δ (ΔUS28). These viruses establish latent and lytic infections, respectively, in CD34^+^ progenitor cells, Kasumi-3 cells, and THP-1 cells ^19,25^, and we confirmed these phenotypes are also maintained in primary CD14^+^ monocytes by supernatant transfer to permissive fibroblasts (Figure 4A). We were also able to detect US28 protein during the establishment of latency in monocytes by immunostaining for the FLAG epitope tag on the C terminus of US28 (Figure 4B).

**Figure 4:**
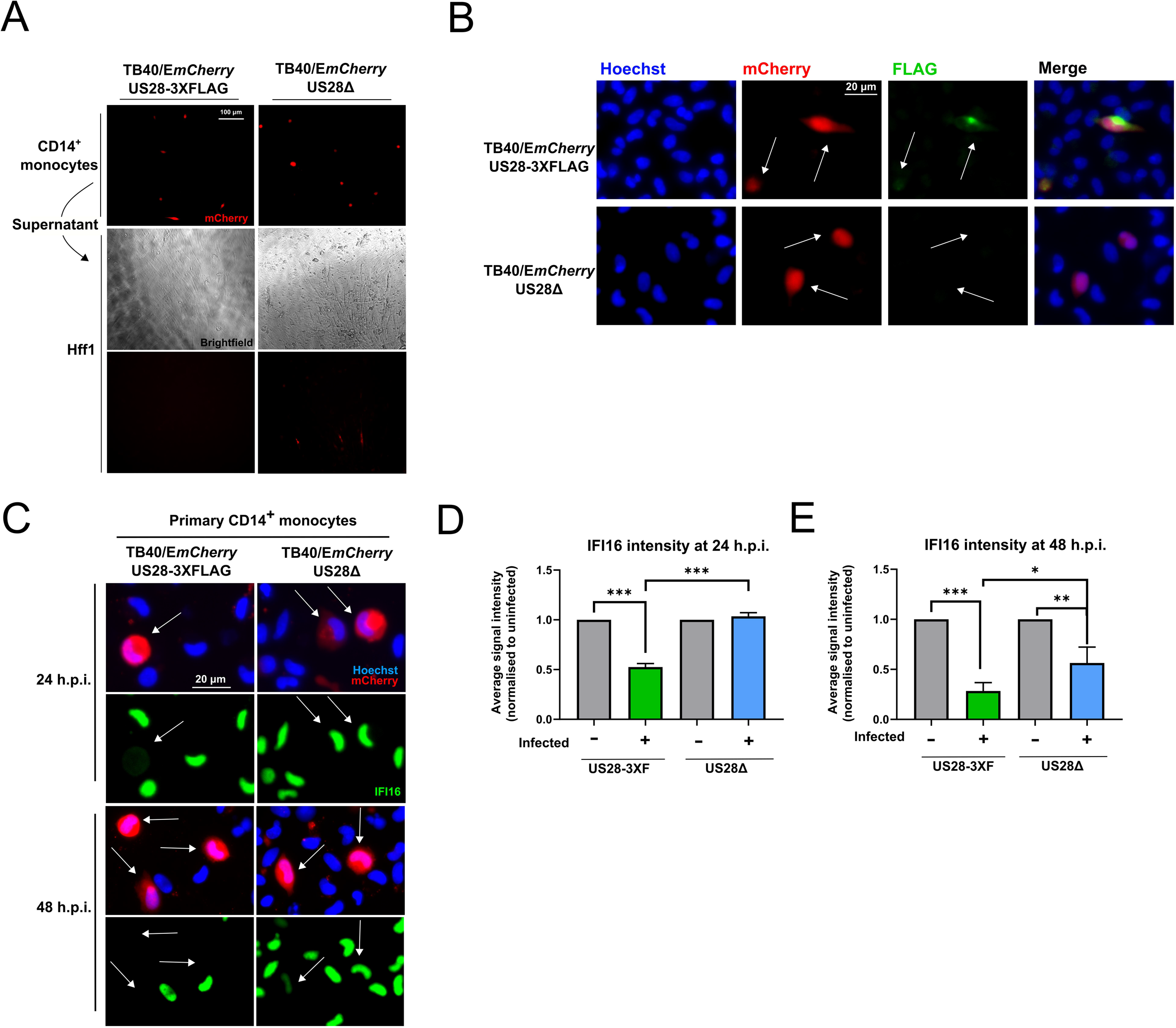
IFI16 is rapidly downregulated in a US28-dependent manner during latent infection. CD14^+^ monocytes were infected with either US28 WT TB40/E*mCherry*-US28-3XFLAG HCMV or the ΔUS28. A) Validation of the latent and lytic phenotypes associated with US28-3xF and ΔUS28 monocyte infections, respectively. At 7 d.p.i., supernatant from infected CD14^+^ cells (upper panel) were transferred to Hff1 cells (middle brightfield and lower mCherry panels) and formation of plaques was monitored and imaged at 20X magnification. B) Detection of US28-3XFLAG during the establishment of latency in CD14^+^ monocytes. At 2 d.p.i. US28-3xF or ΔUS28-infected CD14^+^ monocytes were fixed and stained by immunofluorescence for US28-3XFLAG using an anti-FLAG antibody and imaged at 40X magnification. C) US28-3xF and ΔUS28-infected monocytes were stained by immunofluorescence for IFI16 at the indicated times and imaged using 40X magnification. White arrows indicate corresponding cells. D and E) IFI16 signal intensity in each nucleus was normalised to the average of the uninfected cells in a field of view. The results of three fields of view were then averaged to derive the resulting average signal intensities for each subpopulation of monocytes at the indicated time points infected with US28-3xF or ΔUS28 HCMV. Statistical significance was determined using one-way ANOVA. *** indicates P<0.001, ** indicates P<0.01, and * indicates P<0.05.

To determine if US28 specifically downregulates IFI16 in the context of infection, we compared the expression of this cellular protein in monocytes infected with US28-3XF or ΔUS28. Consistent with Figure 3B, we found that monocytes infected with the US28-3xF virus showed downregulation of IFI16 at 24 and 48 h.p.i., while monocytes infected with ΔUS28 displayed robust IFI16 expression at 24 h.p.i. (Figure 4C and 4D) and only partial downregulation at 48 h.p.i (Figure 4C and 4E). These data demonstrate that the early downregulation of IFI16 in CD14^+^ monocytes is dependent on US28.

### IFI16 is downregulated by US28 only in undifferentiated myeloid cells

We previously showed that US28 modulates cellular signalling pathways in undifferentiated, but not differentiated THP-1 cells ^21^. We were therefore curious as to whether the effects on IFI16 expression were dependent on cellular differentiation status. This is significant because differentiated THP-1 cells and mature dendritic cells are permissive for HCMV lytic infection. To analyse whether these effects are differentiation-dependent, we transduced THP-1 cells with a lentiviral vector that co-expresses US28 and the fluorescent protein Emerald (US28-UbEm), or co-expresses eGFP and Emerald (empty UbEm), as a control. For each population, we sorted the Emerald-positive THP-1 cells by FACS (Figure S4B) and validated US28 expression by RT-qPCR (Figure S4C). We treated half of these cells with PMA in order to induce cellular differentiation. We found that undifferentiated US28-expressing THP-1 cells downregulated IFI16, but PMA-differentiated cells did not downregulate IFI16 (Figure 5A), suggesting only latency-associated expression of US28 attenuates IFI16 expression.

**Figure 5:**
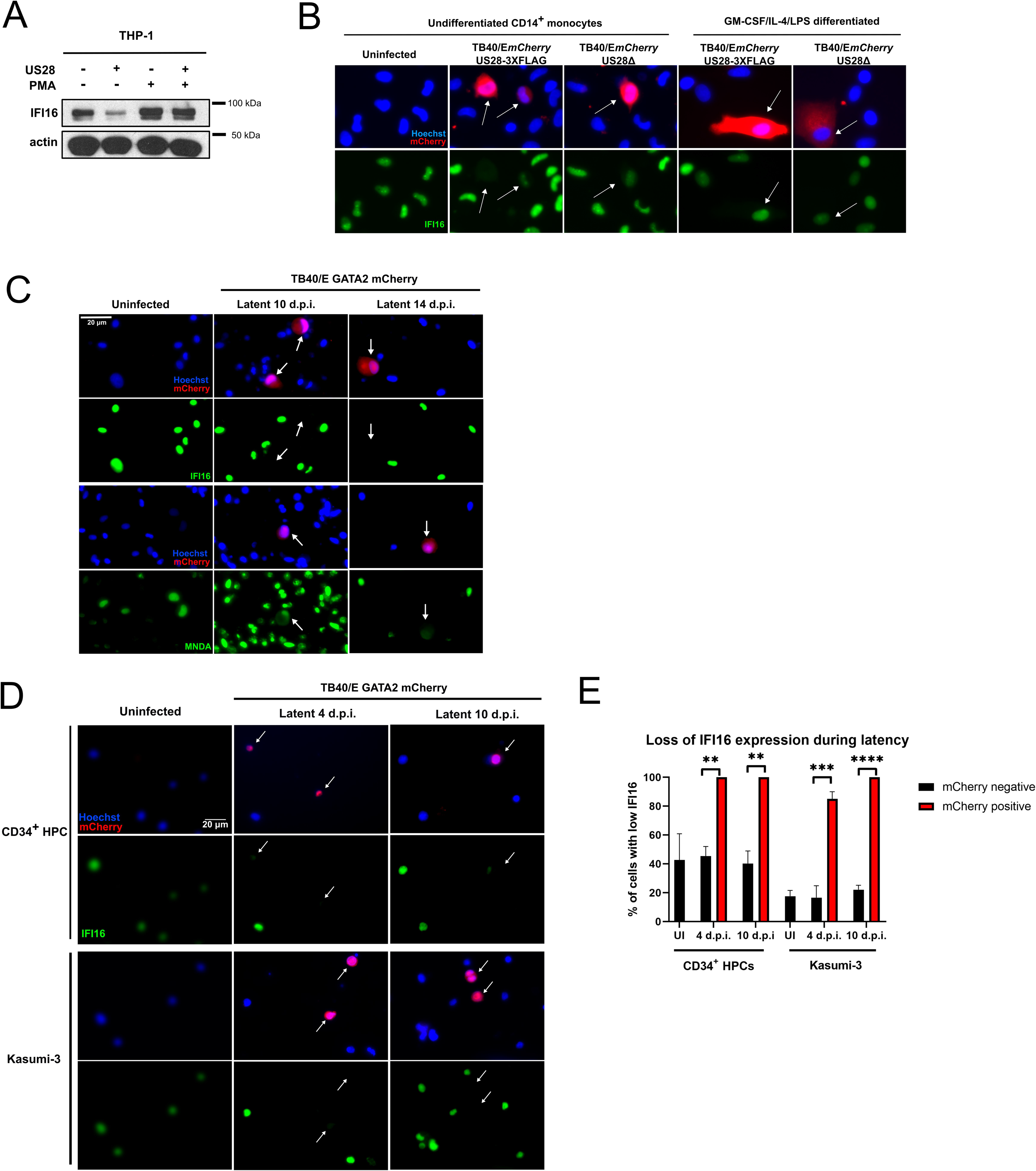
IFI16 downregulation is maintained during long term latency of undifferentiated monocytes and CD34^+^ progenitor cells. A) US28 expressing and empty vector THP-1 cells were either left untreated or treated with PMA for 48 hours before cell lysates were harvested. These lysates were then subject to western blotting for IFI16 and actin as a loading control, with molecular weight markers annotated. B) At 48 h.p.i, either undifferentiated CD14^+^ monocytes, or monocytes pre-differentiated for 7 days with GM-CSF/IL-4/LPS, were fixed and stained for IFI16 and imaged as before at 40X magnification. White arrows indicate corresponding infected cells. C) CD14+ monocytes were infected with HCMV GATA2mCherry, or left uninfected. At the indicated times, cells were fixed and stained for IFI16 or MNDA and imaged as before. D) Primary CD34^+^ hematopoietic progenitor cells from two donors, or Kasumi-3 cells, were infected infected with HCMV GATA2mCherry, or left uninfected. At the indicated times, cells were fixed and stained for IFI16 and imaged as before. E) Quantification of at least 3 fields of view from D), presented as the proportion of cells with low IFI16 expression in the infected, mCherry positive and uninfected, mCherry negative populations. UI – uninfected. Statistical analysis was performed by Fischer’s Exact test on the total number of cells in each category. **** indicates P<0.0001, *** indicates P<0.001, ** indicates P<0.01.

We also analysed the effect of cellular differentiation on IFI16 expression following infection in mature dendritic cells derived by treating *ex vivo* CD14^+^ monocytes with GM-CSF/IL-4/LPS. Again, we found that undifferentiated infected CD14^+^ monocytes downregulate IFI16 in a US28-dependent manner at 48 h.p.i, while infected mature dendritic cells do not downregulate IFI16 with WT or ΔUS28 HCMV (Figure 5B). Taken together, our results indicate that US28 rapidly downregulates IFI16 during latent infection of monocytes, but not during lytic infection of mature dendritic cells.

### Low levels of IFI16 are maintained during long term latency in monocytes and CD34^+^ progenitor cells

We next assessed whether downregulation of IFI16 and MNDA occurs during long term maintenance of latency; long term downregulation of HLA-DR is already known to be important for latent carriage of HCMV^41^. We infected monocytes with HCMV that drives mCherry from GATA2 promoter, and maintains this marker for far longer during latency than SV40 promoter-driven tags^51^. At 10 and 14 d.p.i., IFI16 remained absent and MNDA remained partially downregulated in infected cells (Figure 5C). We then repeated this analysis in primary CD34^+^ hematopoietic progenitor cells (HPCs), a site of long-term *in vivo* latent carriage, as well as the Kasumi-3 cell line, an experimental model for HCMV latency^64^. Consistent with our observations in monocytes, and RNAseq experiments in cord blood derived CD34^+^ cells^65^, IFI16 levels were low or absent in almost all infected cells imaged at 4 and 10 days post infection (Figure 5D and 5E). Thus, it seems likely that downregulation of IFI16 is a conserved process in cellular sites of HCMV latency.

### IFI16 activates IE gene expression via NF-κB in myeloid cells

Having established that US28 downregulates IFI16 early during the establishment of latency, we wanted to address why this may be beneficial to the virus for latent infection. One function of IFI16 is the sensing of viral DNA and subsequent induction of Type I interferon or interleukin-1-beta ^27,65^. While we do not currently rule out a potential role for IFI16-mediated sensing of HCMV during latency, we were more struck by the previous work identifying IFI16 as a modulator of host and viral transcription ^32–34,36,37,66–71^. In particular, IFI16 is capable of activating the MIEP and driving IE gene expression within the first 6 hours of lytic infection of fibroblasts ^32,34^, though at later times IFI16 blocks early and late gene expression ^34,37^. Given that a hallmark of HCMV latency is the suppression of IE gene expression ^72^, we hypothesised that high levels of IFI16 might drive MIEP activity and IE gene expression in undifferentiated myeloid lineage cells. To address this, we transduced and selected THP-1 cells with control empty vector (EV) or IFI16-overexpression lentiviruses to generate control and IFI16-overexpressing cell lines. We validated IFI16 overexpression by western blot (Figure 6A) and then infected these cell lines with recombinant HCMV carrying an IE2-YFP cassette to allow us to identify cells that express IE2^50^. Undifferentiated THP-1 cells are an established model for a number of aspects of HCMV latency^73^, including the significant repression of IE2 expression ^73^. When we infected control and IFI16-overexpressing THP-1 cells with HCMV in five paired experiments, we found a consistent increase in the number of IE positive cells in IFI16-overexpressing THP-1 cells (Figure 6B and 6C), suggesting IFI16 overexpression drives IE protein production in cells that would otherwise repress this viral protein.

**Figure 6:**
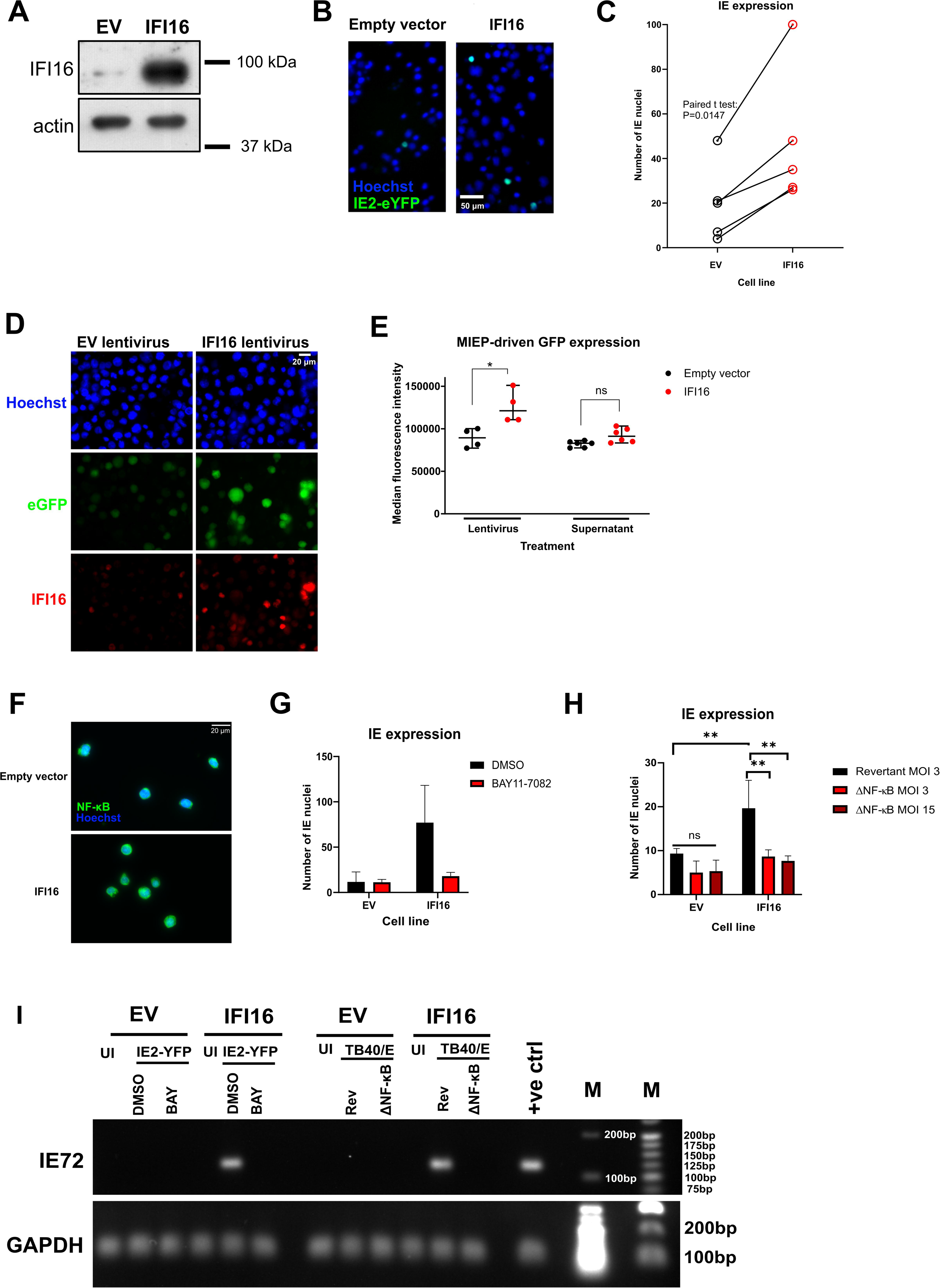
Overexpression of IFI16 in monocytic cells leads to MIEP activation and IE gene expression via NF-κB. A) THP-1 cells were transduced with empty vector (EV) or IFI16-overexpression lentiviruses and following blasticidin selection, IFI16 overexpression was confirmed by western blot. Actin is shown as a loading control. B and C) Empty vector or IFI16-overexpressing cells (from A) were infected with TB40/E IE2-eYFP virus, and IE2-eYFP positive nuclei were imaged and counted by fluorescence microscopy. C) Cumulative results from five paired experiments, which were analysed by paired two-tailed Student’s t-test; * P<0.05. D) EV and IFI16 lentivirus concentration was determined by p24 ELISA (data not shown) and 15 ng p24 equivalents of each lentivirus was used to transduce MIEP-eGFP THP-1 cells. Cells were maintained for two weeks in culture, and IFI16 overexpression was validated by immunofluorescence. E) Left hand comparison: cells described in (D) were assessed for eGFP fluorescence by flow cytometry. Right hand comparison: non-transduced MIEP-eGFP expressing cells were incubated with supernatants from cells described in (A) for two days. eGFP expression was quantified by flow cytometry. A statistical comparison of the median fluorescence intensity was performed using two-tailed Mann-Whitney test; ns, not significant and * P<0.05. F) Empty vector or IFI16-overexpressing cells were fixed and stained for NF-κB, with Hoechst as a nuclear stain, at 40X magnification to assess levels of nuclear NF-κB. G) Empty vector or IFI16-overexpressing cells were infected with TB40/E IE2-eYFP virus in the presence of the IKKα inhibitor BAY11-7082 which inhibits the NF-κB pathway, or DMSO as a control. IE2-eYFP positive nuclei were imaged and counted by fluorescence microscopy at 48 hours post infection. H) Empty vector or IFI16-overexpressing cells were infected with a revertant WT-like TB40/E at MOI 3, TB40/E with NF-κB binding sites deleted from the MIEP (ΔNF-κB) at MOI 3 or MOI 15. At 48 h.p.i., cells were fixed and stained for IE and the number of IE positive nuclei were counted. Graph shows the results of three experiments and statistical analysis by 2-way ANOVA using Sidak’s multiple comparison test. ** P<0.01, ns, P > 0.05. I) Empty vector or IFI16-overexpressing cells were infected as per F) and H) at MOI 3, but cells were instead analysed for IE72 expression by RT-qPCR. PCR products were then run on a 2% (upper panel, IE72) or 1.2% (lower panel, GAPDH) agarose gel. UI refers to uninfected cells, DMSO is the solvent control, BAY refers to BAY11-7082, Rev refers to the revertant TB40/E and ΔNF-κB to the NF-κB binding site mutant virus. The positive control (+ve ctrl) was HCMV-infected PMA-differentiated monocytes. Molecular weight markers (M) are annotated.

Prior work has identified that IFI16 could activate IE gene expression in a UL83-dependent manner during lytic infection ^32,37^, but since this tegument protein does not enter the nucleus in the CD34^+^ progenitor cell model of HCMV latency ^74^, we hypothesised that IFI16 could activate the MIEP without additional virion components. To test this hypothesis, we utilized a THP-1 MIEP reporter system; THP-1 cells in which an integrated 1151 bp region of the MIEP drives the expression of eGFP ^42^. In these undifferentiated THP-1 cells, the MIEP is epigenetically repressed unless stimulated (for example by differentiation) ^42^. We treated these MIEP-eGFP THP-1 cells with control lentiviruses or lentiviruses which drive the overexpression of IFI16, ensuring equivalent lentivirus infection of reporter cells by p24 ELISA. These cultures were maintained for two weeks, after which we validated IFI16 expression by immunofluorescence (Figure 6D) and then analysed eGFP expression by flow cytometry (Figure 6E). We found that the IFI16-overexpressing cells had increased eGFP expression compared with controls (Figure 6D and 6E), suggesting that IFI16 overexpressed in isolation and in the absence of additional HCMV components, drives MIEP activity. Furthermore, culturing THP-1 MIEP-eGFP reporter cells with supernatants from the empty vector or IFI16-overexpressing cell lines in Figure 6A resulted in no significant MIEP activity, suggesting that the effect is mediated intracellularly, and not by a secreted factor (Figure 6E).

IFI16 activates NF-κB signalling in a number of contexts^66,75^, and our previous work indicates that US28-mediated attenuation of NF-κB signalling is important for the establishment of latency^21^. Therefore, we hypothesised that IFI16 activates the MIEP via NF-κB. We found increased nuclear NF-κB localisation in IFI16 overexpressing cells (Figure 6F), and by using the NF-κB pathway inhibitor, BAY11-7082, we were able to ameliorate the effect of IFI16 overexpression on IE (Figure 6G and 6I), suggesting that NF-κB plays an important role in this pathway. Finally, we infected IFI16-overeexpressing cells with HCMV that lacks NF-κB sites within the MIEP^52^ to check whether IFI16 exerts its effects via direct binding of NF-κB to the MIEP. In this case, IFI16 overexpression failed to induce IE gene expression, unlike IFI16 cells infected with the revertant strain (Figure 6H and 6I). Taken together, our data are consistent with the view that IFI16 activates IE gene expression in early myeloid lineage cells by allowing NF-κB to bind at the MIEP.

## Discussion

The viral GPCR US28 is expressed during both lytic and latent infection of HCMV. While US28 is dispensable for lytic replication *in vitro* ^76,77^, it is essential for the establishment and maintenance of HCMV latency in early myeloid lineage cells ^19,21,23,25^. This is attributable, in part, to the ability of US28 to suppress the major immediate early promoter; a US28 function specific for undifferentiated myeloid cells ^18,21,23,25^.

We hypothesised that this ability of US28 to so profoundly regulate viral IE gene expression in undifferentiated myeloid cells was likely via US28-mediated modulation of host protein abundance and, therefore, we wanted to determine whether such US28-driven changes could be important for the establishment or maintenance of HCMV latency. While previous work has used targeted arrays to assess US28-mediated effects on myeloid cells ^21,23,25^; here we have used an unbiased proteomic screen to understand how US28 reprograms host cells in order to support latent infection. Our screen compared host protein abundance in control THP-1 cells or THP-1 cells which express either WT-US28 or the US28 signalling mutant, US28-R129A. As such, we could assess the signalling-dependent and signalling-independent effects of US28. We then chose to focus on signalling-dependent changes because G protein coupling via the residue R129A is essential for experimental latency ^21,25^. However, we predict that some of the signalling-independent changes driven by US28 could also be important for HCMV latency, since these changes included alterations in several cell-surface molecules such as co-stimulatory molecule CD82, adhesion molecule CD44, and in receptor tyrosine kinase FLT3. The latter two cellular factors are implicated in myeloid cell differentiation, which is intimately linked with HCMV latency and reactivation ^18,72,78–80^. As such, modulating these cell-surface molecules could help to control interactions with immune effectors and cellular differentiation-linked reactivation.

By looking at changes in host protein abundance between US28-WT and US28-R129A expressing THP-1 cells, we found a number of significant changes in the host proteome which likely result specifically from US28 signalling. Interestingly, we found US28-WT downregulated a large number of interferon-inducible proteins, including canonical interferon-stimulated genes (ISGs) like OAS2 and IFITM3, as well as MNDA, IFI16, and several HLA-DR components. We found that levels of both total STAT1 and phosphorylated STAT1 were reduced in US28-WT expressing cells, a mechanism that may act in synergy with the US28-mediated attenuation of c-fos to downregulate interferon-inducible genes^25,61^. Modulation of interferon signalling has not previously been reported for US28, but in the context of the latently infected monocyte, a general block in downstream interferon signalling may be important for maintaining the polarisation of the monocyte ^81,82^, or perhaps to avoid the anti-viral activities of ISGs. We believe these questions merit further interrogation.

We chose to focus on the two PYHIN proteins and the set of HLA-DR components which are downregulated by US28. We confirmed downregulation of IFI16, MNDA, and HLA-DR in THP-1 cells which overexpress US28 and recapitulated these effects in experimental latency in primary CD14^+^ monocytes. HLA-DR was previously reported to be downregulated during experimental latency in granulocyte-macrophage progenitor cells, which prevents CD4^+^ T cell recognition and activation ^41,83,84^. Whilst this down-regulation of MHC Class II involved the expression of the latency-associated gene UL111A ^41^, our data argue that viral US28 could also contribute to this phenotype. Little is known about the function of MNDA, a myeloid specific PYHIN protein implicated in neutrophil cell death ^85^ and monocyte transcriptional networks ^86^. Ongoing work in our laboratory aims to identify whether US28-mediated downregulation of MNDA during latent infection could benefit latent carriage and/or reactivation.

Our results clearly characterised a rapid downregulation of IFI16 during the establishment of latency in monocytes, which occurred within the first 24h of infection and was also maintained during long term latency in monocytes and CD34^+^ HPCs. The early downregulation of IFI16 was clearly US28-dependent as ΔUS28 virus failed to display immediate IFI16 down-regulation. However, we did observe a partial downregulation of IFI16 in ΔUS28-infected monocytes at later time points of infection. We think it likely that this involves an unidentified lytic-phase viral gene product, which may be required for overcoming the known IFI16-mediated restriction of HCMV lytic infection ^30,34,36,37^ and occurs as a result of ΔUS28 virus initiating a lytic infection in undifferentiated monocytes.

Our observation that the US28-dependent downregulation of IFI16 occurred rapidly (within 24h of infection) may, in part, be attributable to incoming US28 which is functional ^25^. IFI16 protein has a short half-life of approximately 150 minutes in fibroblasts ^87^ and therefore, incoming US28 protein may rapidly target IFI16 transcription in latently infected monocytes, as it does in US28-expressing THP-1 cells, resulting in loss of IFI16 within 24 hours of infection; this is then maintained by subsequent latency-associated *de novo* US28 expression.

We found that preventing IFI16 expression has a clear benefit to the establishment of HCMV latency. This contrasts with previous analyses of latency in other viral systems, where IFI16 expression is necessary to repress lytic viral transcription ^39,40^. In our study, IFI16 overexpression activated MIEP activity in the absence of additional viral proteins, and, furthermore, IFI16 overexpression increased IE positive nuclei in latently infected THP-1 cells. IFI16 activates the MIEP during lytic infection ^32,34^, though in these cases an additional viral gene product, UL83, is thought to be required. Our results suggest that UL83 is not required for IFI16-mediated activation of the MIEP in undifferentiated myeloid cells, and suggest that IFI16 activates NF-κB to achieve this, as use of either an NF-κB pathway inhibitor or deletion of NF-κB binding sites from the MIEP prevented IFI16-mediated IE expression. We believe this provides one mechanism by which US28 blocks NF-κB activity early during latency, a phenomenon we previously showed to be important for the establishment of latency in myeloid cells^21^.

Taken together, our results suggest that one of the early events in the establishment of latency in CD14^+^ monocytes is the US28-mediated targeting of interferon responsive genes, including the downregulation of IFI16, which serves to support the repression of the MIEP.

## Acknowledgements

We would like to thank Linda Teague, Paula Rayner, Isobel Jarvis, and Roy Whiston for technical assistance and Dr. Ian Groves for critical reading of the manuscript. This work was funded by the British Medical Research Council, Grant (Grant G0701279), the Wellcome Trust (Grant 109075/Z/15/A) and the Cambridge NIHR BRC Cell Phenotyping Hub. The funders had no role in study design, data collection and interpretation, or the decision to submit the work for publication.

## Ethics statement

All human samples were obtained under ethical approval and after approval of protocols from the Cambridgeshire 2 Research Ethics Committee (REC reference 97/092) and all protocols were conducted in accordance with the Declaration of Helsinki. Informed written consent was obtained from all of the volunteers included in this study prior to providing blood samples and all experiments were carried out in accordance with the approved guidelines.

## Author contributions

E.G.E., B.K., J.W., E.Y.L, G.X.S., P.J.L., and J.S. designed experiments. E.G.E., B.K., J.W., E.Y.L., E.P., G.X.S., and J.S. performed experiments and data analysis. E.G.E. and J.W. wrote the manuscript. C.M.O., M.W., P.J.L, J.S. supervised the research. E.G.E., B.K., J.W., E.Y.L, M.W., C.M.O., P.J.L., and J.S. edited the manuscript.

---

**Figure S1. US28 expression induces IFN-inducible genes, but not ER stress-related genes**

A) Changes in interferon inducible genes identified in Figure 1D, and other canonical ISGs, in US28-WT with respect to US28-R129A. Green bars indicate changes with a q value of <0.001. D) Heat map of the changes in canonical ER stress-related genes induced by US28-WT or US28-R129A expression as per the proteomic screens in Figure 1A, B, C. HUGO gene symbols are listed followed by a common gene name, if applicable. An outgroup of genes that are regulated by US28 (IFI16, MNDA, FLT3) is included for comparison.

**Figure S2. US28-expressing cell lines downregulate IFI16, MNDA, and HLA-DR**

A) Empty vector, US28-WT and US28-R129A-expressing THP-1 cells were regenerated in independent transductions using the same expression vectors as used for the proteomic screen (Figure 1). US28 expression was validated by RT-qPCR, with US28 RNA normalised to TATA-box binding protein (TBP) and presented as 2^−ΔCt^. B) Cells from A were lysed and subject to western blot for US28, and actin as a loading control. C) Quantification of three western blots for US28 expression. C and D) Lysates prepared from cells in (A) were analysed by western blot for IFI16 (C) and MNDA (D) expression; actin is shown as a loading control. Note that panel E is from the same membrane as Figure 1C. F) Quantification of 5 and 4 independent western blots for IFI16 and MNDA, respectively. G) Cells from A) were treated with ruxolitinib as per figure 2D, or left untreated. Lysates from these cells were analysed by western blot for phosphorylated STAT1, total STAT1, or actin as a loading control.

**Figure S3. Strain-dependent differences in US28 do not affect downregulation of interferon-inducible genes**

A) Sequences encoding US28 from the indicated HCMV strains or plasmids were aligned using Clustal Omega. B) Retroviral plasmids encoding US28-WT (from TB40/E) or R129A, each with a C-terminal 3XFLAG tag, and an eGFP marker, were used to transduce THP-1 cells. They were then subject to immunofluorescence staining for the 3XFLAG tag. C and D) Cells from B were stained for cell-surface HLA-DR by flow cytometry. D) Mean fluorescence intensity of the US28-WT and US28-R129A cell lines. Statistical analysis by Student’s t test; ** P<0.01.

**Figure S4 Downregulation of IFI16, MNDA, and HLA-DR is not simply a bystander effect of contact with viral particles.**

A) CD14^+^ monocytes were left uninfected, or infected with HCMV for 24 hours before fixing and staining for the indicated proteins, and imaging as before. B) The sequence encoding US28 from VHL/E was cloned into the lentiviral plasmid pUbEm (US28-UbEm), and this or empty UbEm plasmid was used to transduce THP-1 cells, which were subsequently cell-sorted for Emerald expression. C) US28 expression was validated in the cells from (B) by RT-qPCR. US28 RNA was normalised to cellular TBP and presented as 2^−ΔCt^.

**File S5: US28 proteome in THP-1 cells**

Tab 1: Data: THP-1 cells expressing empty vector, US28-WT, and US28-R129A were subject to total cell proteomic analysis using a TMT labelling approach as described in Materials and Methods. This file lists all genes identified in this proteomic screen, including their Uniprot Accession number, HUGO gene symbol, fold changes in abundance between cell lines, and q values of statistical significance. Tab 2: Interferome Top 40 Downreg. Gene names, fold changes, and q values of the top 40-most downregulated genes (US28-WT vs US28-R129A) are presented, along with whether they are included in the Interferome database as being Type I or Type II interferon-inducible (marked with ‘y’). Tab 3: Interferome Zero-Change 40: Gene names, fold changes, and q values of the genes with a fold change value of zero (US28-WT vs US28-R129A) are presented, along with whether they are included in the Interferome database as being Type I or Type II interferon-inducible (marked with ‘y’).

**File S6: Schematic showing mass spectrometry settings for experiments presented in Figure 1 and File S5**

