## Supplementary figures and images for "Interferon-responsive genes are targeted during the establishment of human cytomegalovirus latency"

### Figure S1

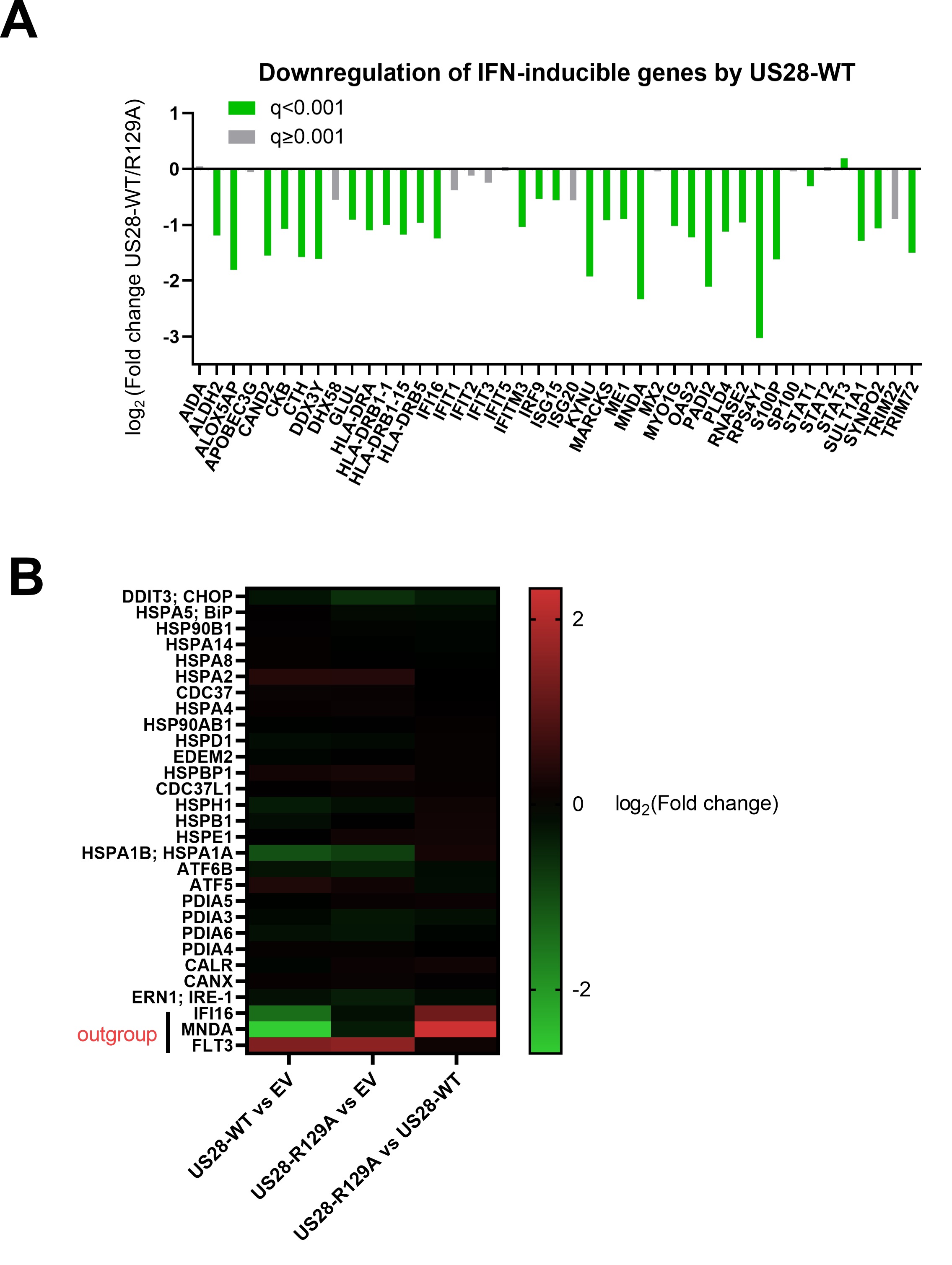

### Figure S2

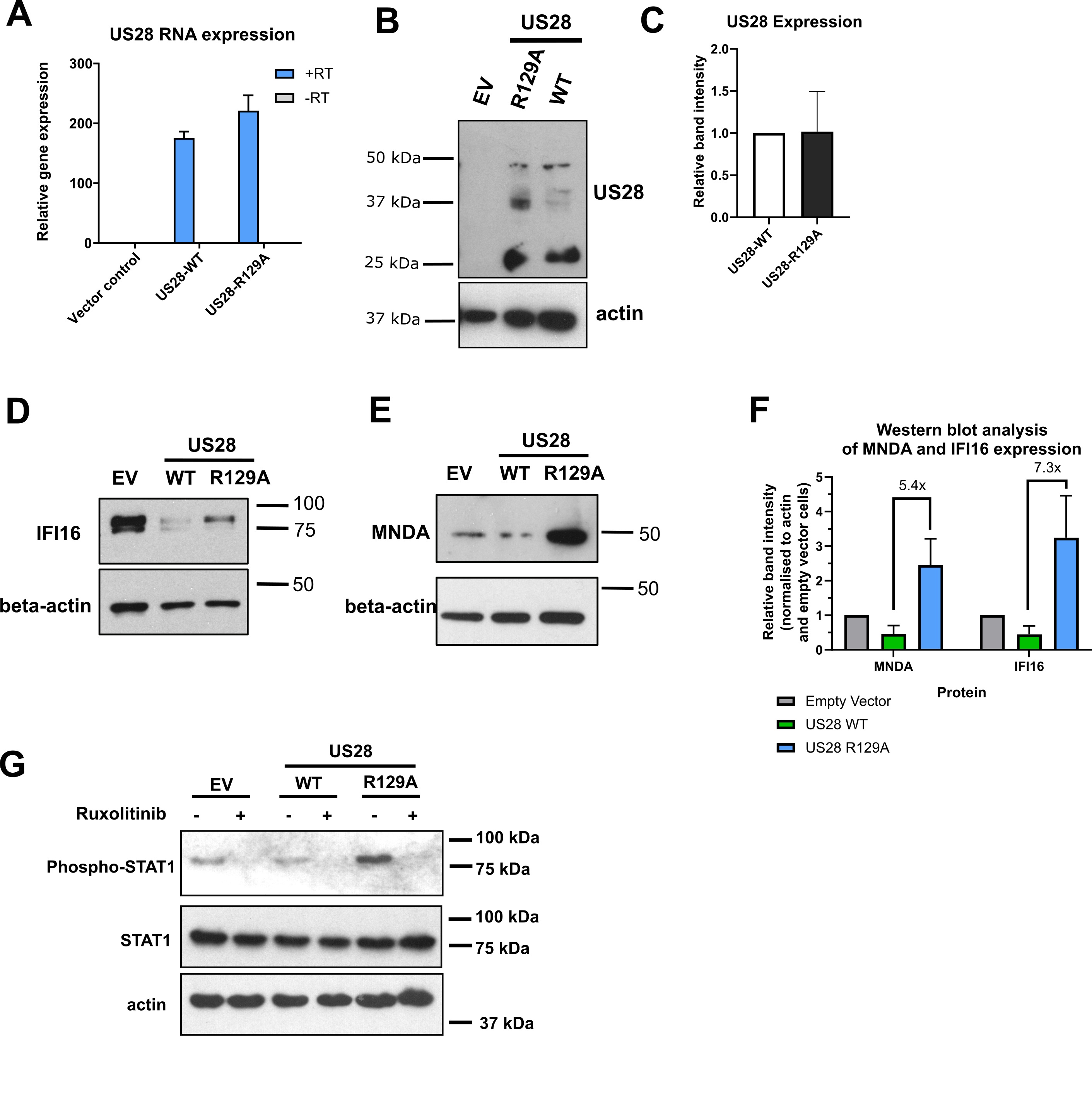

### Figure S3

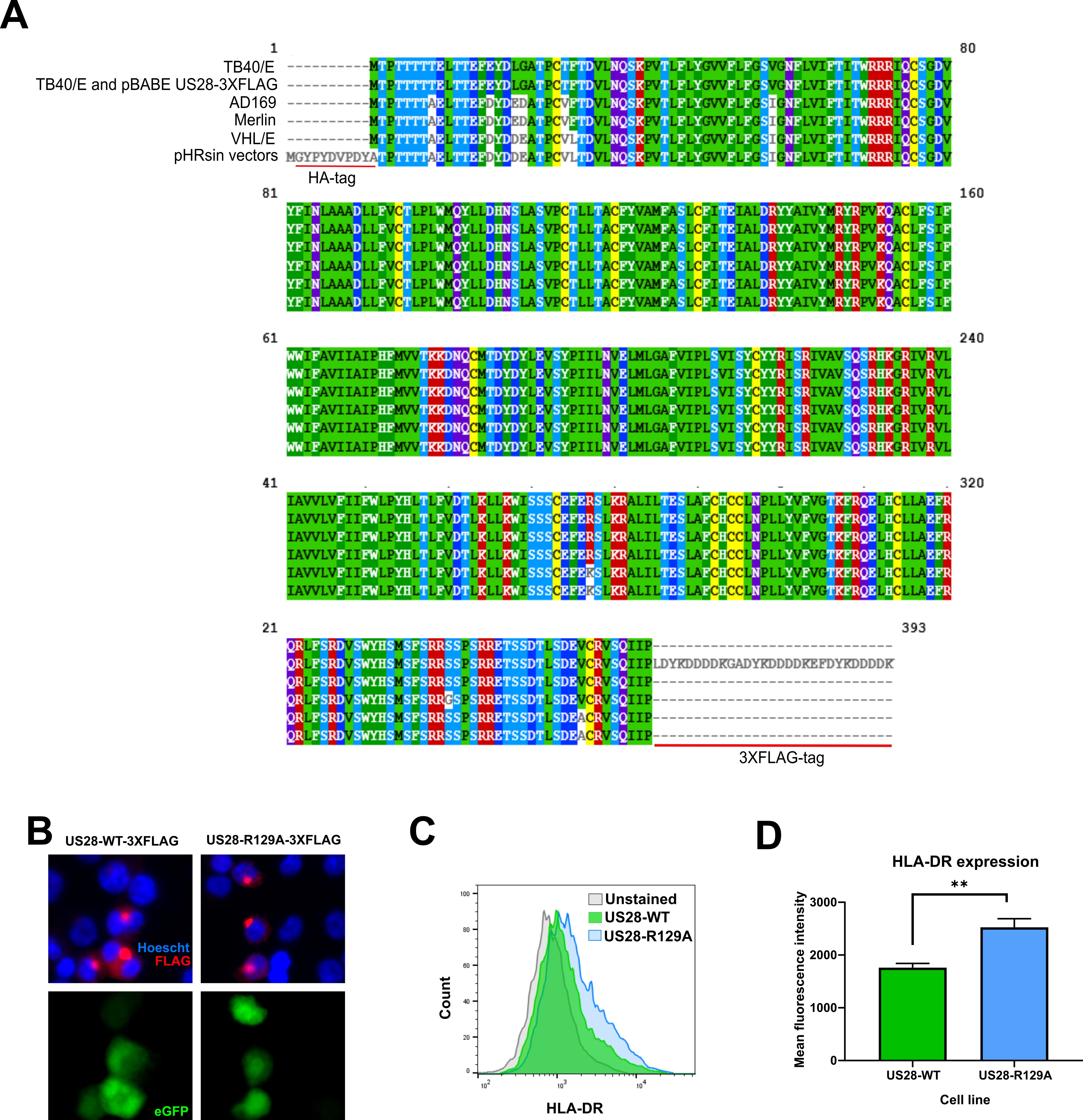

### Figure S4

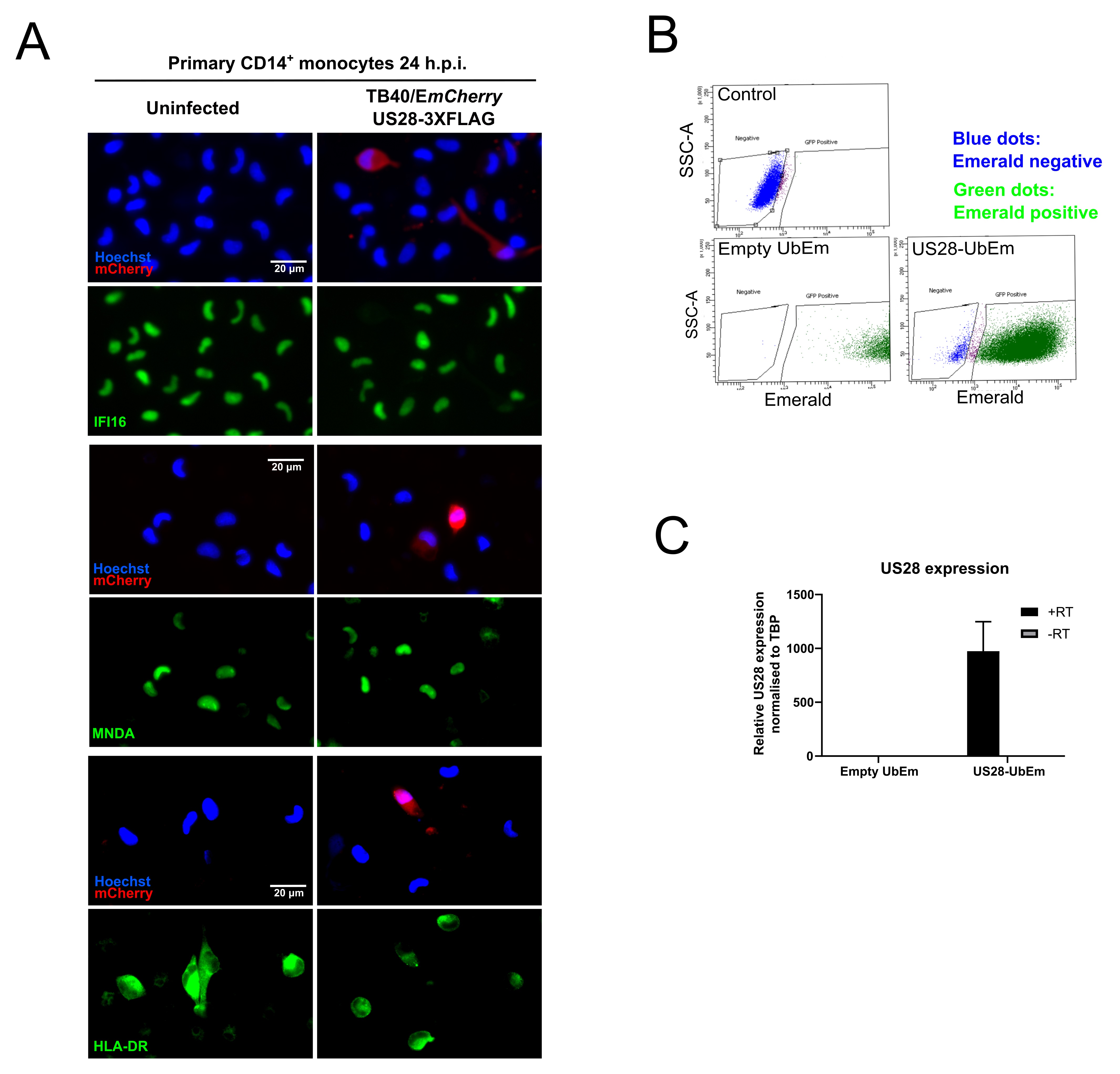
